# Development and validation of a computational tool to predict treatment outcomes in cells from High-Grade Serous Ovarian Cancer patients

**DOI:** 10.1101/2024.10.02.616212

**Authors:** Marilisa Cortesi, Dongli Liu, Elyse Powell, Ellen Barlow, Kristina Warton, Emanuele Giordano, Caroline E. Ford

**Author notes:** Corresponding author(s). E-mail(s),; Contributing authors.

## Abstract

Treatment of High-Grade Serous Ovarian Cancer (HGSOC) is often ineffective due to frequent late-stage diagnosis and development of resistance to therapy. Timely selection of the most effective (combination of) drug(s) for each patient would improve outcomes, however the tools currently available to clinicians are poorly suited to the task.

We here present a computational simulator capable of recapitulating cell response to treatment in ovarian cancer. The technical development of the in silico framework is described, together with its validation on both cell lines and patient-derived laboratory models. A calibration procedure to identify the parameters that best recapitulate each patient’s response is also presented.

Our results support the use of this tool in preclinical research, to provide relevant insights into HGSOC behaviour and progression. They also provide a proof of concept for its use as a personalised medicine tool and support disease monitoring and treatment selection.

## 1 Introduction

High-grade serous ovarian cancer (HGSOC) is the most common subtype of epithelial ovarian cancer, accounting for approximately 70% of cases [1]. The silent nature of this disease, which often lacks specifically recognisable symptoms at its early stages, leads to most patients (about 85%) being diagnosed after it has spread beyond the reproductive organs [2]. This complicates the clinical management of HGSOC since late diagnosis increases the rates of recurrence and treatment resistance [3] and reduces 5-year survival to less than 50% [4].

Treatment of HGSOC mostly relies on debulking surgery and combined administration of platinum and taxane agents. In homologous-recombination deficient (HRD) patients, PARP-inhibitors are also used as a maintenance therapy to induce synthetic lethality [5]. Upon relapse, other chemotherapeutic agents, anti-angiogenics, immunotherapy and targeted treatments can be introduced [6]. Despite the increasing number of options, treatment effectiveness in HGSOC remains low and the identification of the best drug for each patient remains a significant challenge. While the limited number of established markers of response is a major hindrance to patient stratification, disease heterogeneity is emerging as the main issue. Indeed, HGSOC is characterised by a quick evolution, which often leads to several genetically distinct sub-clones and different drug resistance mechanisms coexisting within the same patient [7].

We here explore the use of digital twins, which are computational models recapitulating features specific to individual patients such as individualised response to treatment, to address these limitations. Digital twins have been successfully applied to treatment personalisation in several conditions [8–12], but no HGSOC-specific tool is currently available. We chose to focus on recreating a virtual representation of the lining of the peritoneum (the omentum), as HGSOC primarily spreads within the abdominal cavity (transcoelomic metastasis). The tool that we have created, named ALISON (digitAl twIn Simulator Ovarian caNcer), mimics the metastatic process by combining the simulation of individual cells (healthy and cancer) using an agent-based framework, with the finite element modelling of the concentration distributions of key molecules (oxygen, glucose, lactate, drugs) and their passive diffusion within the virtual tissue. The inclusion of features like cell heterogeneity and the interaction between cancer and healthy cells increase the *in vivo* relevance of this framework and of its results.

The technical development of ALISON will be described in this paper, together with its experimental validation in cultured cell lines and patient-derived samples. A digital twin calibration procedure is also introduced, to enable the identification of the parameters best recapitulating each patient’s response to treatment from clinical data.

Our results support the use of ALISON in the pre-clinical setting, to complement experimental data and study HGSOC behaviour and progression. They also suggest the feasibility of ALISON as a personalised medicine tool, capable of supporting disease monitoring and treatment selection. The programmable, modular nature of ALISON also enables its extension to other diseases, using this work as a roadmap for development and testing.

## 2 Results

### 2.1 ALISON development

ALISON combines an agent-based framework (AB) and a finite element model (FEM). The former describes the behaviour of individual cells through a set of programmable probabilistic rules, while the latter simulates the diffusion of key molecules (oxygen, glucose, lactate, drugs) and their distribution within the virtual tissue. Both models rely on the same 3D mesh and extensively interact throughout the simulation. Cells consume oxygen and glucose and produce lactate, thus altering the concentration distributions of these molecules, which in turn affects cell behaviour.

ALISON’s base structure is a schematic representation of the omentum. It comprises a base of fibroblasts and extracellular matrix, topped by a dense layer of mesothelial cells that constitutes the interface with the abdominal cavity (Figure 1a). HGSOC cells can be present in the peritoneal space (uppermost region of Figure 1a) where the pathological accumulation of fluid (ascites) facilitates cancer cell dissemination and metastasis formation [13, 14]. The mesh used in our simulations (Figure 1b) replicates this layered structure, with size and dimensions set so as to be coherent with those of the organotypic model of the omentum [15] which was used as the experimental cell culture model for most of the analysis. It is important to note that the three layers in Figure 1b outline the regions where virtual cells were initially seeded. Throughout the simulation, cancer cells can migrate downwards taking the place of mesothelial cells that have died, or displacing them as they colonise the healthy tissue.

**Fig. 1.**
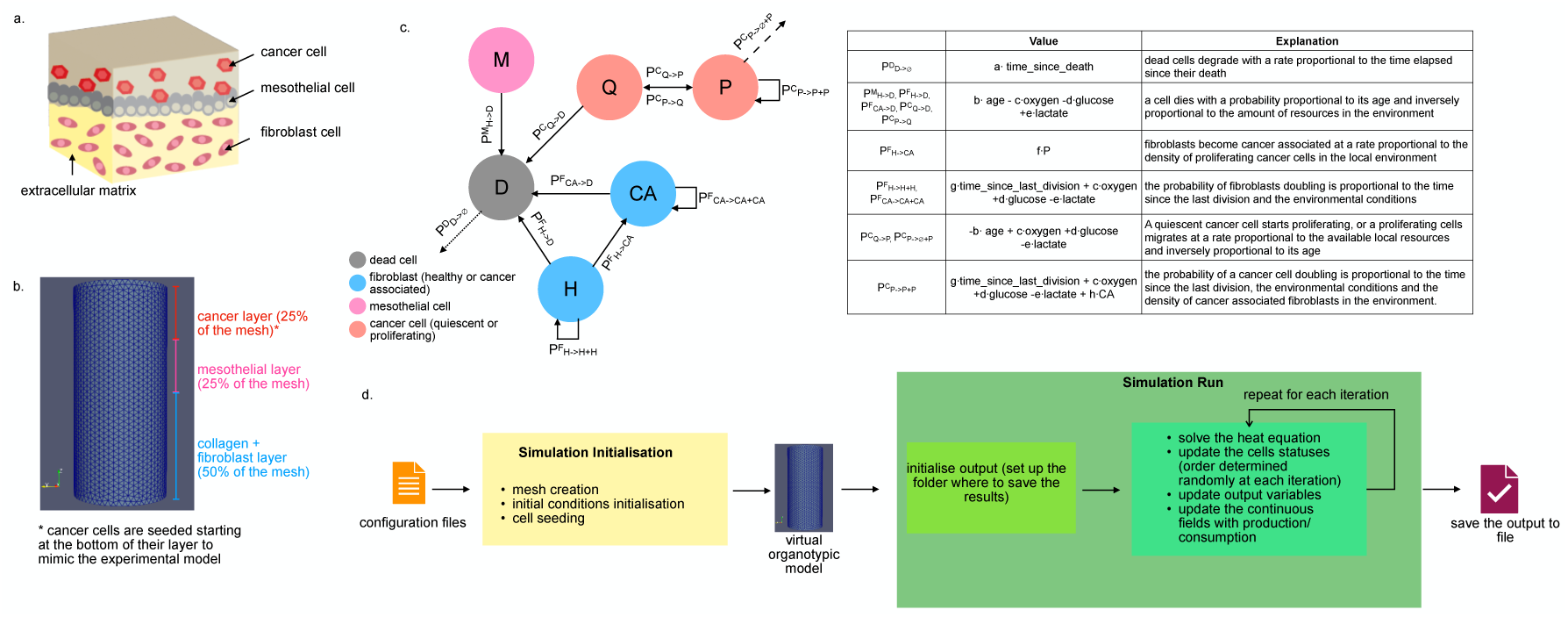
Outline of ALISON and its simulation. a. Schematic structure of the omentum tissue. A base of fibroblasts embedded in the extracellular matrix is overlaid by a continuous layer of mesothelial cells, which constitute the interface between the omentum and the peritoneal cavity. Cancer cells are present in this region since transcoelomic metastasis is the main dissemination mechanism for HGSOC. b. Mesh used as a base for the model. Bars on the side outline the three layers implemented in ALISON to mimic the structure of the organotypic model of the omentum. c. Structure of the AB model outlining the different cell statuses, the allowed state transitions and the corresponding probabilities of execution (Table). d. Schematic representation of the main steps of an ALISON simulation. Configuration files provide the information necessary to initialise the simulation and construct the virtual organotypic model. The simulation then proceeds by setting up the output folder and repeating the steps in the green box on the right a number of times equal to the number of iterations. The output is then saved to file for further elaboration.

Cancer cell migration and all the other behaviours formalised in the AB model are summarised in Figure 1c. Four cell types (cancer, fibroblast, mesothelial and dead) are considered. Nodes in the graph in Figure 1c represent the different states that the virtual cells can assume, while arrows indicate the allowed transitions between states. Each state transition is characterised by a probability of occurrence (Figure 1c) which varies with cell-specific features (e.g., age, time since last division), environmental conditions (e.g., local levels of oxygen, glucose and lactate) or the number of specific cell types in the microenvironment (e.g., the number of cancer-associated fibroblasts). Each rule is also characterised by a series of parameters (a-h) which are used to weigh the contribution of the different variables. These parameters were identified empirically, by comparing simulated and experimental results, as further described in the following sections.

Each ALISON simulation proceeds as shown in Figure 1d. The input of the simulator is a series of configuration files (examples are provided, together with the code for the simulator, at https://github.com/MarilisaCortesi/ALISONsimulator). These are used to define the behaviour of each cell type, the structure of the experimental model (e.g., the layers of the organotypic model) and the experimental conditions.

This information is used to initialise the mesh, place the cells and set the initial levels of oxygen, glucose, lactate and drug. Then, for each iteration, the distributions of the continuous variables are updated to account for passive diffusion (solution of the heat equation) and one rule for each cell is executed. The update order for the cells is randomly determined at each iteration and the rule to execute is chosen according to their probability. All cell types can also maintain the current status with a probability equal to the complement of all the others. At the end of the simulation, the results are automatically saved. The position and type of each cell, together with the distributions of oxygen, glucose, lactate and drugs are saved for each time point.

This effectively creates a single framework in which cells consume oxygen and glucose and produce lactate at realistic rates (Table A1), and the likelihood of the formalised cell behaviours depends on environmental variables (e.g., the local level of oxygen or glucose), similar to that previously described by our team [16–18].

### 2.2 Identification of parameters

The behaviour of virtual organotypic models containing only fibroblasts or mesothelial cells was initially simulated by changing the parameters between 0 and 1.

The number of living cells at the end of each simulation was computed and compared with the starting condition. As limited information on the behaviour of healthy cells in the organotypic model was available, we therefore assumed their density to be constant within the experiments. As such, the parameter sets associated with the smallest difference between initial and final cell density were selected (Table A2).

A similar approach was applied to the identification of the parameters for the panel of six HGSOC cell lines (Table 1) initially used for ALISON’s validation. In this case, the organotypic model comprising all three cell types was simulated and the results were compared to the experimentally measured doubling rates (Table 1), the adhesion dynamics and invasion rates. A cost function further detailed in the methods was used to measure the concordance between each simulated configuration and the experimental data to determine the accuracy of each parameter set in replicating each cell line’s in vitro behaviour. The resulting optimal parameters are available in Table A3.

**Table 1.** Summary of the cell line panel used in this work. Cell lines were chosen as their features resemble those of a HGSOC patient cohort, mostly late-stage with a fair proportion of drug-resistant phenotypes.

| Cell line | Origin | Features | Platinum Response | Doubling time [h] |
| --- | --- | --- | --- | --- |
| CaOV3 | ovarian tissue | First diagnosis, BRCA1/2 wild type, no other known mutations. | sensitive | 78 [39] |
| OAW28 | ascites | Recurrent disease, BRCA1/2 wild type, no other known mutations | resistant | 60 [40] |
| OVCAR4 | ascites | BRCA1/2 wild type | resistant | 43 [41] |
| OVSAHO | abdominal metastasis | BRCA1/2 wild type | sensitive | 56 [42] |
| PEO1 | ascites | First recurrence BRCA2 mutated | sensitive | 37 [43] |
| PEO4 | ascites | Second recurrence (same patient as PEO1). BRCA1/2 wild type following reversion mutation, mutation in NF1 | resistant | 36 [43] |

The experimental quantification of adhesion was characterised by a high variability among the replicates, often coupled with a tendency for the optical density to increase with the incubation time (Figure 2a). This is consistent with a progressive adhesion of cancer cells onto the substrate and is coherent with other reports from the literature [19, 20].

**Fig. 2.**
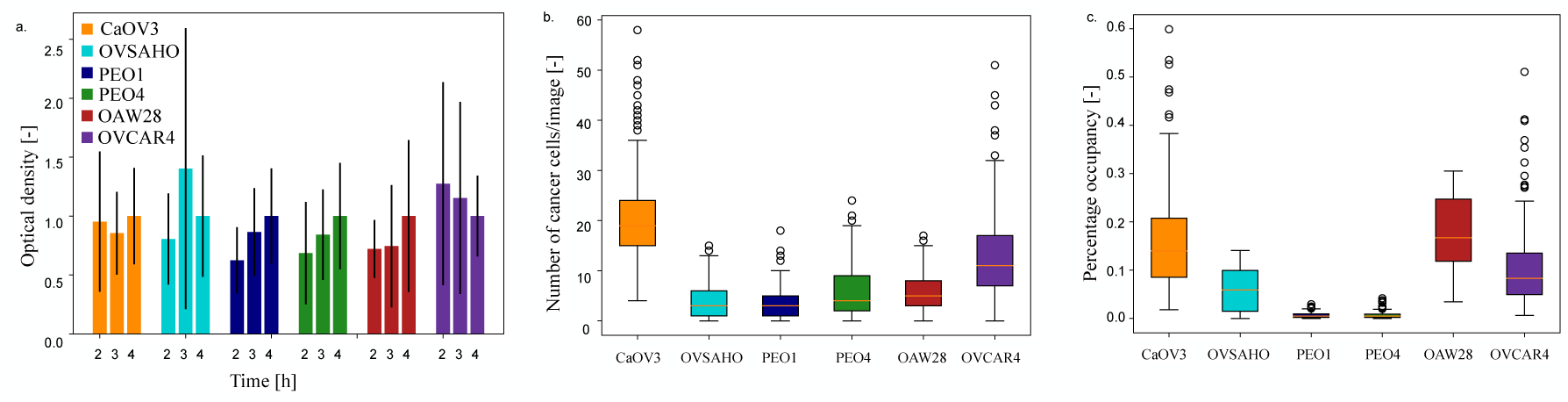
Experimental data measured on the cell lines panel for ALISON’s validation. a. Adhesion timecourse measured at 2, 3 and 4 h after seeding on the organotypic model. Results are shown as average +/- SD of at least 3 independent experiments. b. Number of invading cancer cells measured during transwell invasion assays using ORACLE [21]. Results are shown as average +/- SD of at least 3 independent experiments. c. Percentage of image area occupied by the cancer cells.

Figure 2b and c focus on cell invasion. Transwell assays were conducted using the organotypic model to test the ability of each cell line to invade through this surrogate tissue. The number of invading cancer cells per image was automatically counted using ORACLE [21], a software tool that we developed to distinguish HGSOC cancer cells from healthy cells in co-culture images. The median number of cancer cells/image (Figure 2b) varies between 5 and 20, with a notable variation in distribution width among the considered cell lines. Most of the recognised cells are classified as cancer (Table 2), with an average fraction of cancer cells above 80% for 4 out of the 6 cell lines. OVSAHO and PEO1 cells were associated with the highest fraction of healthy cells and also the lowest number of cancer cells/image (Figure 2b and Table 2). Interestingly, both these cell lines were derived from a drug-sensitive metastatic tumour (Table 1) suggesting that this disease stage might be characterised by a comparatively low cancer cell migration but an increased ability to affect the environment and the behaviour of the healthy cells therein.

**Table 2.** Average fraction of cells recognised as cancer during the analysis. Data reported as average +/- SD.

| Cell line | Fraction of cancer cells |
| --- | --- |
| CaOV3 | 0.995 +/- 0.021 |
| OVSAHO | 0.766 +/- 0.416 |
| PEO1 | 0.692 +/- 0.411 |
| PEO4 | 0.824 +/- 0.342 |
| OAW28 | 0.951 +/- 0.200 |
| OVCAR4 | 0.907 +/- 0.213 |

While further analysis would be required to confirm this hypothesis, this is consistent with the increased plasticity of metastatic cancer cells, which have been shown to remodel their environment to better support their growth [22].

Figure 2c, finally, shows how the fraction of the image area occupied by the cancer cells varies across the different cell lines. This was used as a proxy for the cancer cell prevalence within the healthy tissue, which is the metric used to evaluate the invasion potential in our simulations. Similarly to Figure 2b, the area occupied by CaOV3 cells is quite high, then it decreases in the following 3 cell lines. OAW28 and OVCAR4 cells, on the other hand, occupy an area comparable to CaOV3 cells.

This suggests a non-linear dependence between invasion and disease progression, with highly aggressive early and late-stage diseases (CaOV3, OAW28, OVCAR4) separated by a relatively low motility phase (OVSAHO, PEO1 and PEO4), possibly connected with the establishment of new lesions and their molecular evolution. This intermediate stage might also be characterised by a more extensive interaction between cancer and healthy cells (see the higher fraction of migrated healthy cells in Table 2). These results highlight the value of complex *in vitro* experimental models and their analysis methods in enabling the study of the interactions among different cell types and how they affect cell behaviour.

### 2.3 ALISON validation in HGSOC cell lines

ALISON’s validation in HGSOC cell lines involved simulating dose-response curves to cisplatin, carboplatin and paclitaxel, and comparing them to experimental measurements acquired in comparable conditions. For these simulations, all virtual cancer cells were assigned the set of parameters best representing the corresponding cell line, that is the one characterised by the lowest score, according to the analysis outlined in the previous section. Figure 3 shows the results of this analysis for all drugs and cell lines. Simulated and experimental data were largely consistent, with similar overall dynamics and changes in viability mostly coherent between simulated and experimental data.

**Fig. 3.**
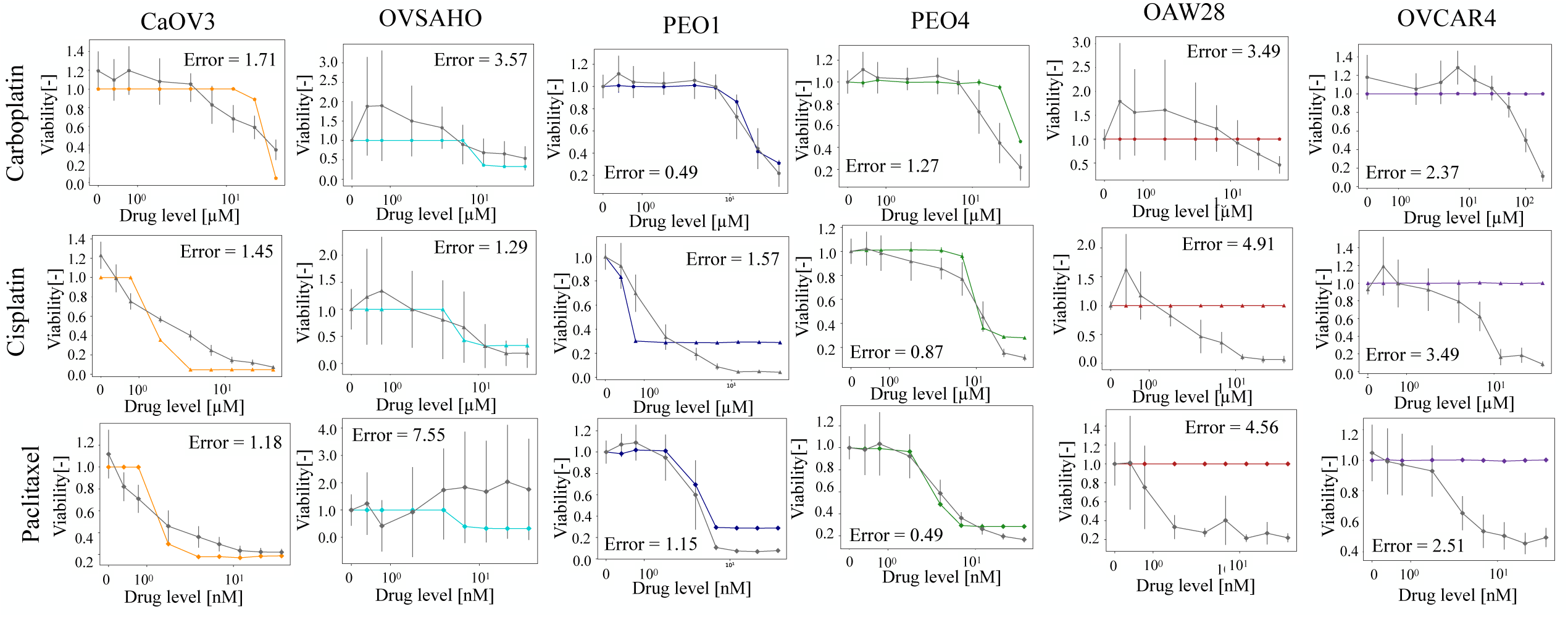
Comparison between the simulated and measured response to carboplatin, cisplatin and paclitaxel. Simulated tracks are colour-coded according to the cell line they refer to, while grey identifies the experimental data. Results are shown as average +/- SD. 10 simulations and 3 independent *in vitro* measurements were considered. Error computed using Equation 1.

The error function in Equation 1 was used to quantify the difference between simulated and experimental data. It measures the overall difference in viability between *in silico* and *in vitro* results at all tested drug concentrations (d). The absolute value is also used to avoid differences in opposite directions cancelling each other out. Low errors are indicative of good quantitative agreement between simulated and measured viability while higher values of this metric correspond to a larger discrepancy.

Cell lines representing an advanced disease stage (OAW28 and OVCAR4) are characterised by a higher error. Indeed, the simulation of these cell lines shows no response to treatment (flat red and purple lines in Figure 3), while a partial to moderate response is observed experimentally, especially at the higher drug doses (grey tracks in the same panels of Figure 3). These results suggest that ALISON might be magnifying each cell line’s characteristics, with drug-resistant ones being completely unresponsive to treatment and early-stage models showing a tendency to respond more quickly.

Comparing the experimentally measured *IC*_50_ with the corresponding value obtained from the simulations further confirms these considerations (Figure 4a). Indeed, most cell lines and drugs show a good correlation, with the major discrepancies being observed for OAW28 and OVCAR4. The Pearson’s correlation coefficient shows a dependence on the cell line and drug treatment (Figure 4b) with carboplatin treatment and earlier-stage disease cell lines being the most accurately captured. Cisplatin and paclitaxel treatments are associated with the lowest correlation. This result is entirely due to the discrepancy observed for OAW28 and OVCAR4 cells. Removing these cell lines from the calculation leads to a remarkable increase in correlation coefficients (0.98 and 0.92 respectively). Overall, the correlation between simulated and experimental *IC*_50_ is quite high (*R*^2^ = 0.85) and consistent with an accurate recapitulation of experimental behaviour by our model.

**Fig. 4.**
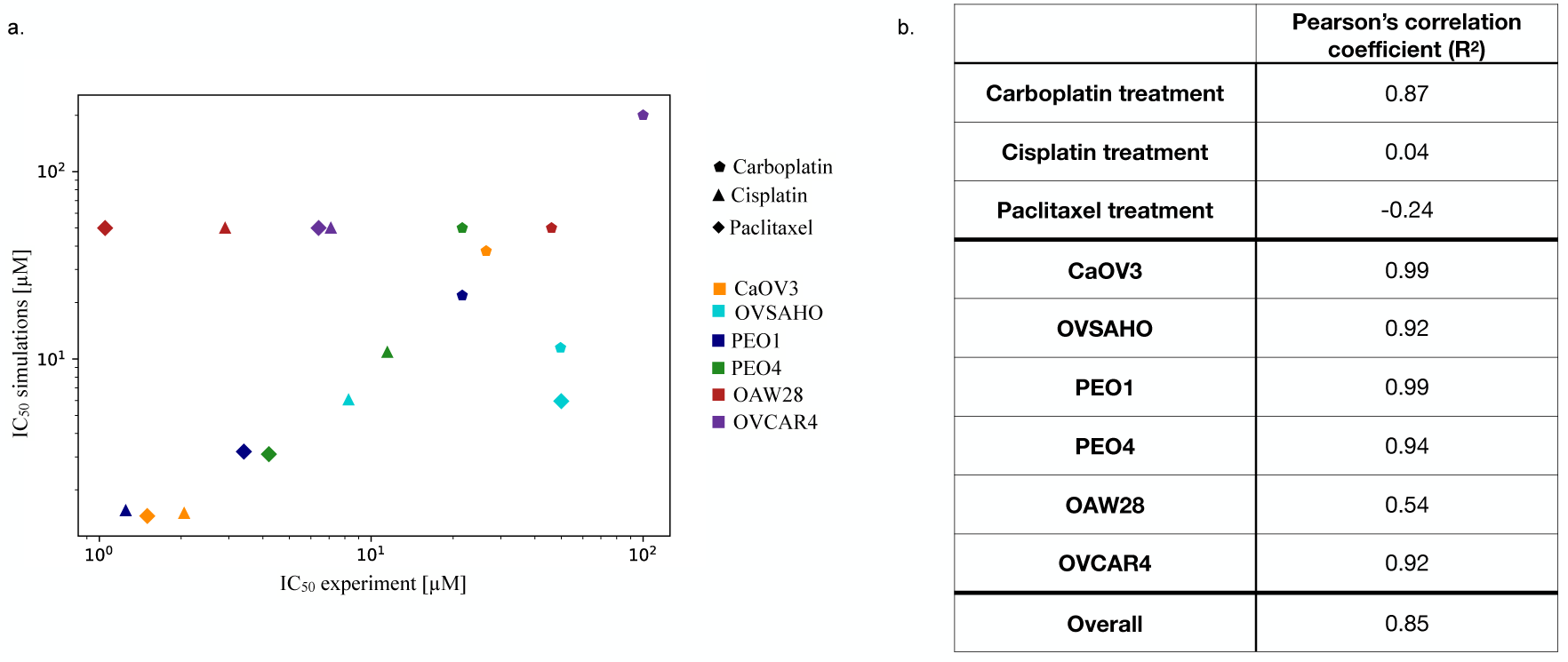
a. Scatter plot analysing the correlation between experimental and simulated *IC*50. Colours identify cell lines while marker shapes are associated with different drugs. b. Correlation coefficients obtained using all available data (Overall) or by grouping them according to cell lines or drugs.

### 2.4 Simulation of cell heterogeneity

Cell heterogeneity has a fundamental role in cancer progression and metastasis, having been connected with the development of treatment resistance, and the ability to withstand harsh, variable environmental conditions [23–26].

Since HGSOC is characterised by a significant diversity both among individuals and within the same patient, we decided to adapt the populations of models approach [27] to our application. This method consists of assigning each cell a different parameter set, chosen according to experimentally relevant criteria. Single-cell level *in vitro* measurements are generally used, but we relied on the distribution of score values calculated during the parameter identification step to establish a proportionality between the likelihood of choosing a parameter configuration and its accuracy in mimicking the behaviour of the corresponding cell line.

The analysis shown in Figure 3 was then repeated to evaluate how cell heterogeneity would affect the outcome of the simulations (Figure 5). Directly comparing the simulated curves in Figure 5 with those in Figure 3 highlighted an interesting behaviour. Cell lines representative of a later disease stage (OAW28, OVCAR4) are better captured by the simulations including cell heterogeneity (Figure A1) while the error for the other cell lines tends to increase with an inverse proportionality to the disease stage (Figure A1).

**Fig. 5.**
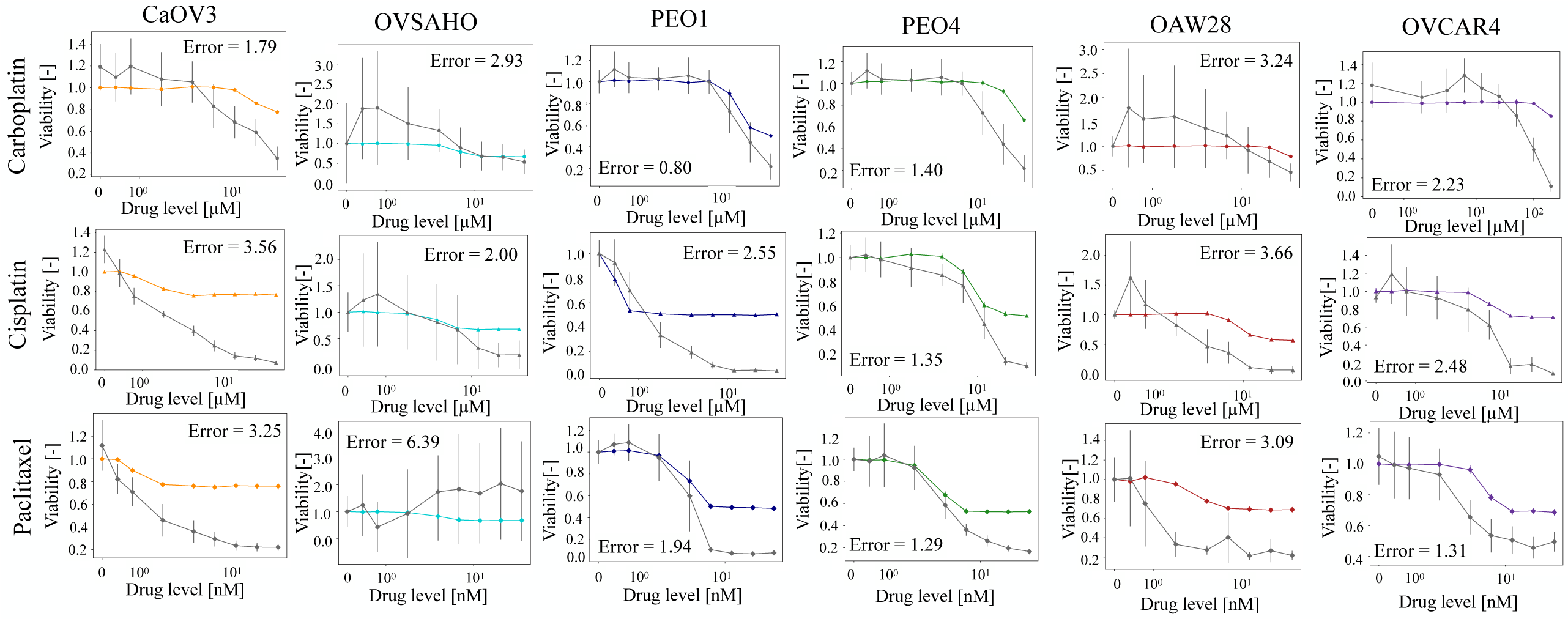
Comparison between the simulated and measured response to carboplatin, cisplatin and paclitaxel when cell-cell variability was included. Simulated tracks are colour-coded according to the cell line they refer to while grey identifies the experimental data. Results are shown as average +/- SD. 10 simulations and 3 independent in vitro measurements were considered. Error is computed using Equation 1.

This result suggests that the simulation of cell-cell variability can mimic the fundamental role of population heterogeneity in HGSOC progression, with an approximately uniform ovarian tumour, that gradually evolves to be more heterogeneous as it spreads beyond its original site and acquires drug resistance [7, 28].

The comparison between the simulated and experimental *IC*_50_s also shows an improvement in correlation and agreement (Figure 6a). A higher *R*^2^ is measured for all three treatments, while cell lines either maintain their correlation value or increase it (Figure 6b). The only exception is PEO4 cells which show a decrease in correlation of about 6%.

**Fig. 6.**
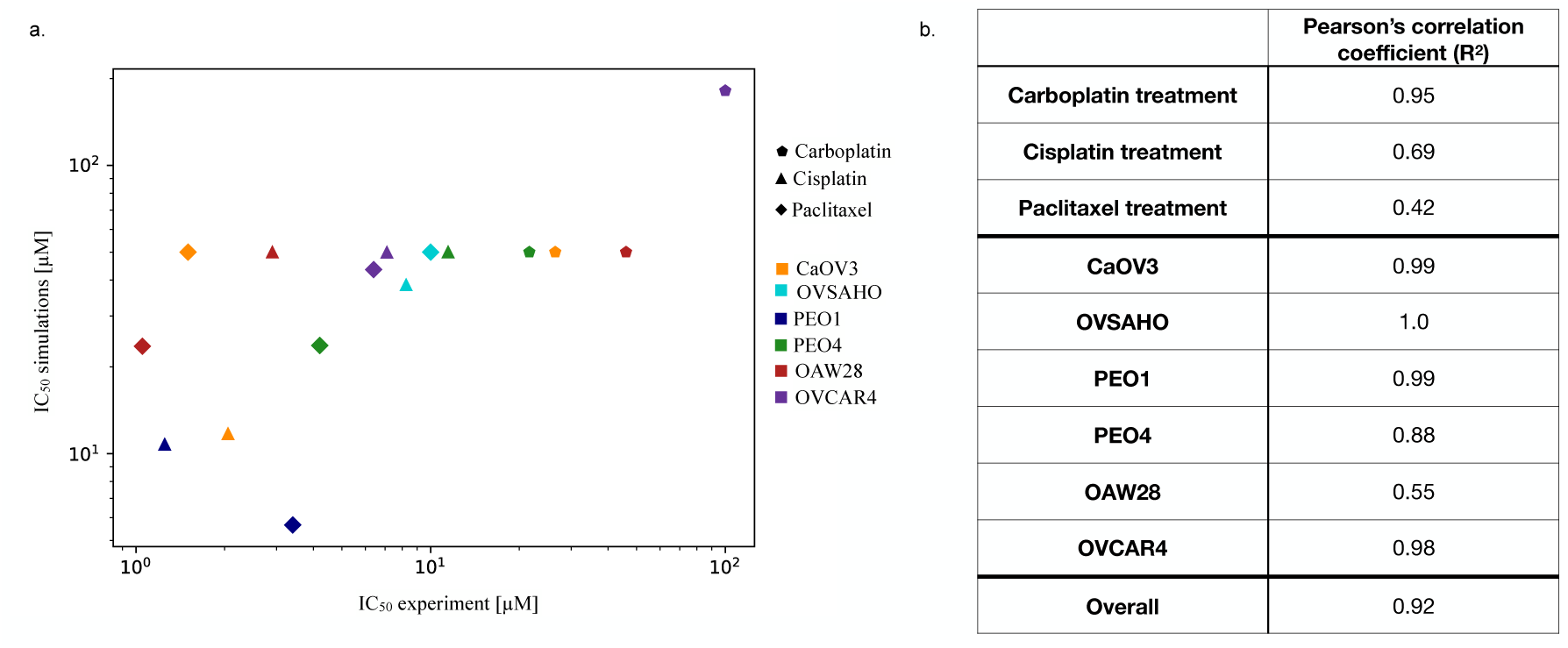
a. Scatter plot analysing the correlation between experimental and simulated *IC*50 in the presence of cell-cell variability. Colours identify cell lines while marker shapes are associated with different drugs. b. Correlation coefficients obtained using all available data (Overall) or by grouping them according to cell lines or drugs.

**Fig. 7.**
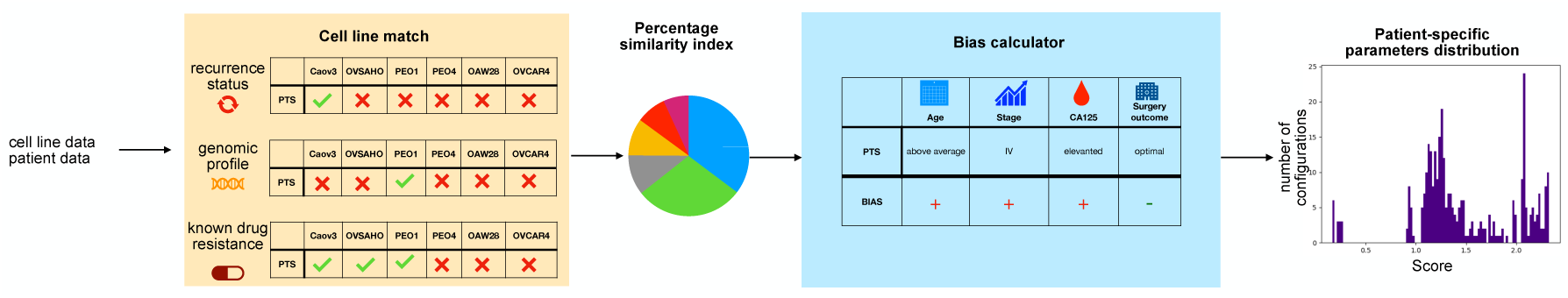
Schematic representation of the digital twin calibration procedure. Recurrence status (e.g. first diagnosis or 2nd recurrence), genomic profile (e.g. BRCA1/2 mutations) and known drug resistance data were matched to the same features in the cell lines panel to determine which combination of cell lines best represents each patient (Percentage similarity index). A prognosis bias was then calculated. It takes into account the age of the patient, the disease stage, the level of CA125, a tumour marker commonly used to diagnose/monitor HGSOC, and the surgery outcome (e.g. if there was residual disease). The overall bias, obtained by adding each individual component, is used to adjust the percentage similarity index. These adjusted fractions are then used as weights to create the patient-specific parameters distribution.

Simulations have a bias toward a worse therapeutic response (higher *IC*_50_) which is consistent with the effect of increased heterogeneity but also provides a more accurate overall estimation of treatment response, as demonstrated by the increase in overall correlation coefficient (*R*^2^ = 0.92).

### 2.5 Digital twins calibration

Following the validation in HGSOC cell lines the digital twins of six HGSOC patients (Table 3) were calibrated according to the procedure schematically represented in Figure 8.

**Fig. 8.**
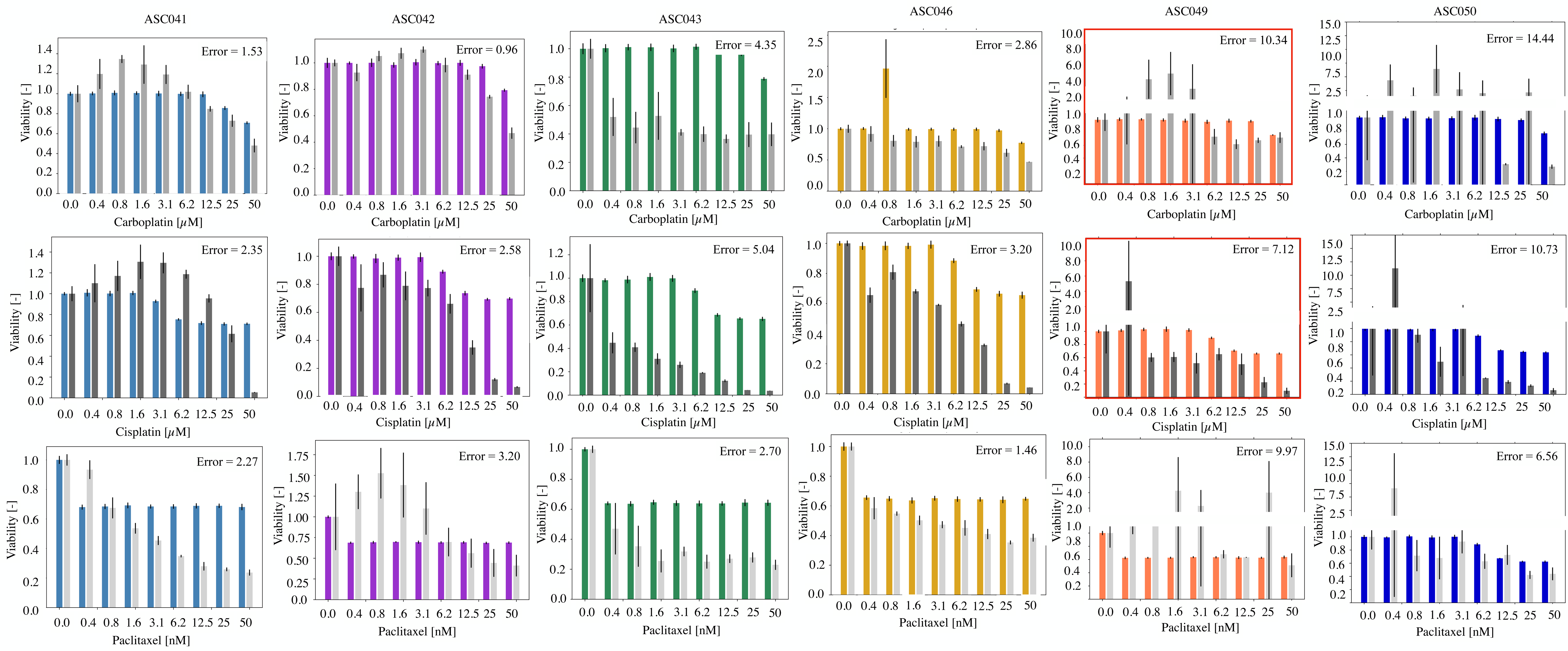
Comparison between experimental and simulated data for ascites-derived cancer cells. Each column represents a patient while rows are associated with different drugs. Coloured bars are the simulated results while grey ones show the experimental data. Results are reported as average +/- SD. 10 simulations and 3 independent in vitro measurements were considered. Error is computed using Equation 1. Red outlines treatments that the patient was known to be resistant to.

**Fig. 9.**
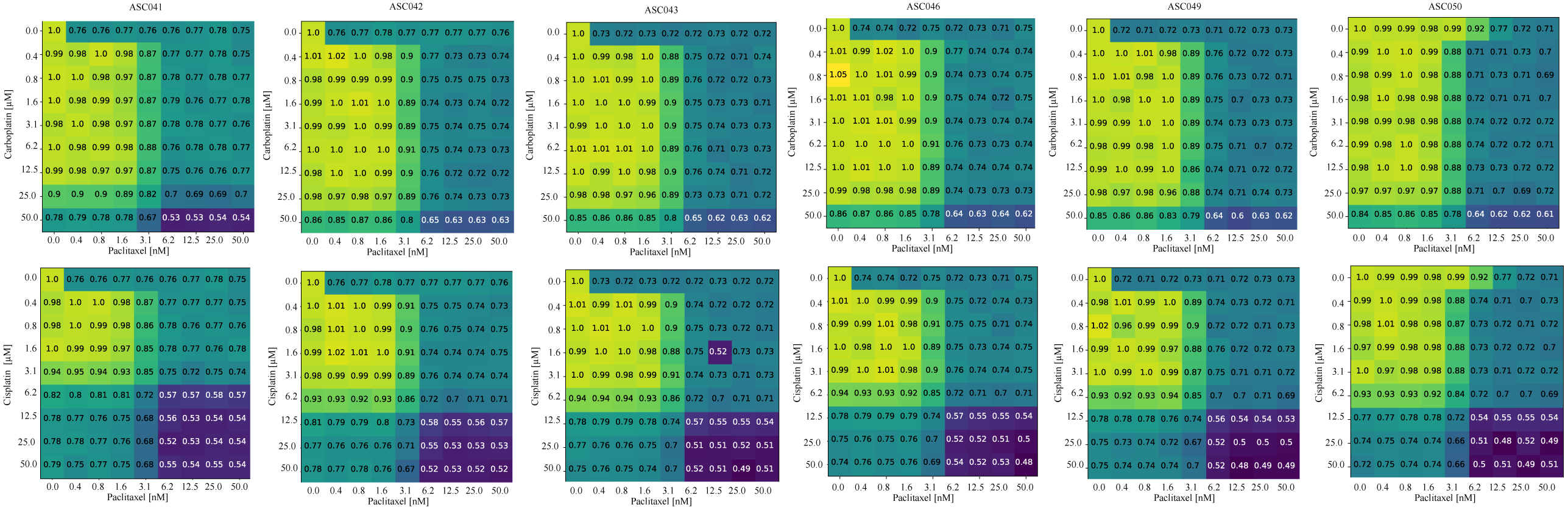
Simulated analysis of the combined treatment with cisplatin/carboplatin and paclitaxel on the digital twins. The numbers and colours represent the average reduction in cell viability with respect to the untreated control. A total of 10 simulations for each condition were considered.

**Table 3.**
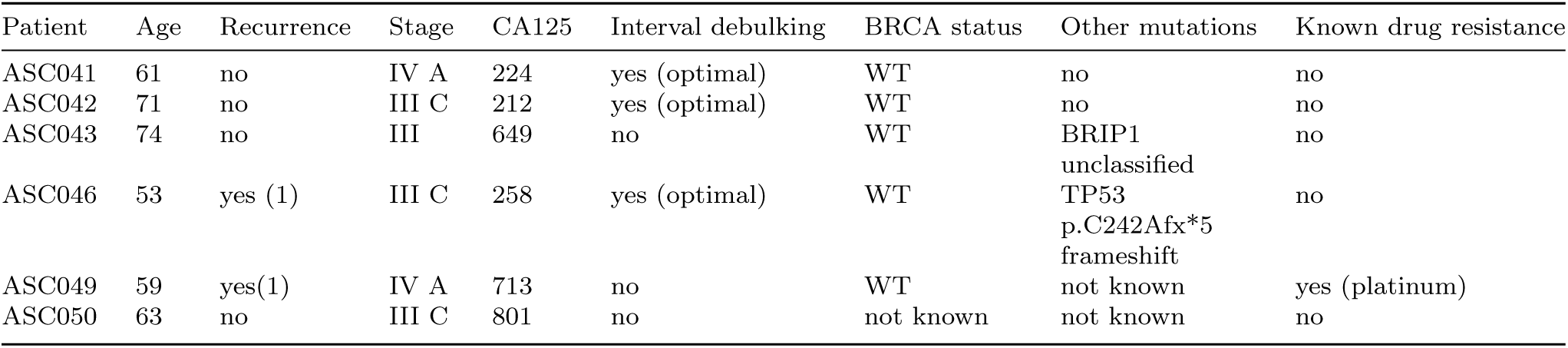
Characteristic of the patients in our cohort.

Initially, the recurrence status, genomic profile and known drug resistance data were matched to the corresponding cell lines’ values to calculate a percentage similarity index, that is the combination of cell lines most accurately describing each patient’s features. These unbiased fractions are reported in Table 4.

**Table 4.** Digital twins calibration procedure. The unbiased fractions are the result of the first step of the procedure, where cell lines and patient features are matched. Biased fractions, on the other hand, are the results of the second stage, where specific clinical features were used to calculate a bias (Table A4) and skew the unbiased fractions toward a better or worse prognosis.

|  | Cell line | ASC041 | ASC042 | ASC043 | ASC046 | ASC49s | ASC050 |
| --- | --- | --- | --- | --- | --- | --- | --- |
| Unbiased fractions | CaOV3 | 0.0 | 0.0 | 0.0 | 0.06 | 0.19 | 0.0 |
|  | OVSAHO | 0.0 | 0.0 | 0.0 | 0.0 | 0.19 | 0.0 |
|  | PEO1 | 0.15 | 0.15 | 0.15 | 0.11 | 0.26 | 0.07 |
|  | PEO4 | 0.50 | 0.50 | 0.50 | 0.50 | 0.26 | 0.29 |
|  | OAW28 | 0.15 | 0.15 | 0.15 | 0.11 | 0.0 | 0.21 |
|  | OVCAR4 | 0.20 | 0.20 | 0.20 | 0.22 | 0.10 | 0.43 |
| Biased fractions | CaOV3 | 0.0 | 0.0 | 0.0 | 0.01 | 0.0 | 0.0 |
|  | OVSAHO | 0.0 | 0.0 | 0.0 | 0.0 | 0.0 | 0.0 |
|  | PEO1 | 0.09 | 0.03 | 0.0 | 0.03 | 0.0 | 0.0 |
|  | PEO4 | 0.50 | 0.37 | 0.39 | 0.38 | 0.36 | 0.33 |
|  | OAW28 | 0.18 | 0.30 | 0.30 | 0.27 | 0.31 | 0.31 |
|  | OVCAR4 | 0.22 | 0.31 | 0.31 | 0.30 | 0.33 | 0.36 |

A prognosis bias taking into account the remaining clinical information (age, stage, CA125 level and interval debulking) was then obtained (Table A4). A negative bias would be associated with a favourable prognosis and thus increase the prevalence of earlier-stage HGSOC models (CaOV3, OVSAHO, PEO1). A positive bias, conversely, would skew the combination toward cell lines representative of a later disease stage (PEO4, OAW28, OVCAR4). The final, biased cell lines fractions are reported in Table 4 (biased fractions), where colours indicate the direction of change. Blue identifies cell lines whose prevalence was decreased while orange those that were increased. All the patients in our cohort had a positive bias, mostly due to having a late-stage disease and an elevated CA125. This led, in the final combination, to a limited representation of early-stage cell lines, whose prevalence in the final fractions was reduced in favour of OAW28 and OVCAR4.

The weighted average of the cell lines’ scores, using the biased fractions as weights, was then calculated to construct the score distribution for each patient.

### 2.6 Digital twins simulation

The response to carboplatin, cisplatin and paclitaxel for each patient was simulated using the digital twins (Figure 8). In all cases, a limited to moderate response to each drug is observed in the simulations. This is mostly coherent with the experimental data acquired on the ascites-derived cells collected from the same patients (grey bars). The error between simulated and experimental data was calculated using the same metric used for the cell lines panel (Equation 1). Overall, a wider range of errors is observed for primary cells, from accurately recapitulated drug responses with low error (e.g., carboplatin for ASC042) to low accuracy configurations (e.g. ASC046 cisplatin). Interestingly, some complex, highly variable responses (e.g. ASC049 carboplatin) are captured fairly accurately by the model but are nevertheless associated with a high error due to the definition of the metric.

Treatment with cisplatin tended to elicit a larger change in cell viability in our *in vitro* experiments, when compared with the simulations, especially at higher drug concentrations. This was also the case for patient ASC049, whose cancer was known to be platinum-resistant (red outlines in Figure 8). The difference between the cisplatin concentration used in the clinic (maximum plasma concentration of 14.4 *µ*M according to [**?**]) and the range considered for the experiments (up to 50 *µ*M) is likely responsible for this discrepancy. Considering a cisplatin concentration of 12.5 *µ*M, which is the closest to the clinically relevant value, patient ASC049 showed a response comparable to that measured for 0.8 *µ*M of the same drug, suggesting a resistant phenotype. Excluding the last two points in the calculation of the error for the cisplatin treatment leads to a reduction in this metric between 10 and 47% thus strengthening the performance of ALISON at clinically relevant drug doses.

The average error across all treatments is the highest for patients ASC049 and ASC050. In both cases, the experimental data show a non-monotonic response to the considered drugs which the simulations are unable to capture. Modifications to the sigmoidal drug response model implemented in ALISON might be required to recapitulate these more complex dynamics.

Patient ASC043 also exhibits a more extensive response to platinum agents, when compared to their digital twin simulation. The *in silico* response is coherent with the biassed fractions in Table 4 which are associated with a more aggressive and treatment-resistant phenotype. A different weighting of the considered patient features might be required to distinguish between the *in vitro* response of patient ASC043 and ASC046. Both of these patients, however, passed away within less than a year from sample collection, suggesting that the presented *in vitro* data might not accurately capture the observed clinical response.

### 2.7 Combined treatment in the digital twins

Standard treatment for HGSOC combines a platinum agent and paclitaxel. We simulated this treatment considering each possible combination of the concentrations used in the previous analysis. Figure 8 shows the average decrease in cell viability, normalised with respect to the untreated control. In all cases, four distinct areas are noted. A top left region (yellow-green squares) where the paclitaxel and/or platinum concentration is low and a limited decrease in viability is observed. A bottom-right corner (dark blue region), where high concentrations of both drugs are associated with the highest decrease in cell population, especially pronounced when cisplatin is considered. Teal-coloured areas in the upper right and lower left corners are associated with an intermediated response which is comparable with single-agent treatment. The size of these regions is fairly consistent across patients, as their response to these combined treatments is fairly uniform. A correlation with the biassed fractions in Table 4 can however be observed. Patient ASC041 is the most responsive with a ratio between the yellow-green and blue area equal to 0.11 and 0.64 for the carboplatin-paclitaxel and cisplatin-paclitaxel treatments respectively. Patient ASC050, on the other hand, exhibits the most drug-resistant phenotype with the same ratios dropping to 0.1 and 0.39. The rest of the cohort is characterised by an intermediate behaviour (Table 5). An analysis of the simulation variability was then conducted by calculating, for each condition, the squared coefficient of variation at the end of the simulation (Equation 2).

**Table 5.** Ratio of the yellow-green (viability 0.8) and dark blue (viability 0.65) for the combined treatments in. **Figure 8**.

| Patient | Yellow-green to blue area ratio |  |
| --- | --- | --- |
|  | Carboplatin + paclitaxel | Cisplatin + paclitaxel |
| ASC041 | 0.11 | 0.64 |
| ASC042 | 0.10 | 0.43 |
| ASC043 | 0.10 | 0.50 |
| ASC046 | 0.10 | 0.46 |
| ASC049 | 0.10 | 0.46 |
| ASC050 | 0.10 | 0.39 |

Figure 10 shows the results of this analysis. Two main patterns of inter-simulation variability were observed. Most patients tended to exhibit higher variability when high doses of both paclitaxel and the platinum-based agent were administered (patients ASC041, ASC046, ASC049, ASC050). In platinum-resistant and generally low-response patients (ASC046, ASC049, ASC050) this trend is less pronounced with occasional high variability configurations at low drug doses, but it is still clearly observable.

**Fig. 10.**
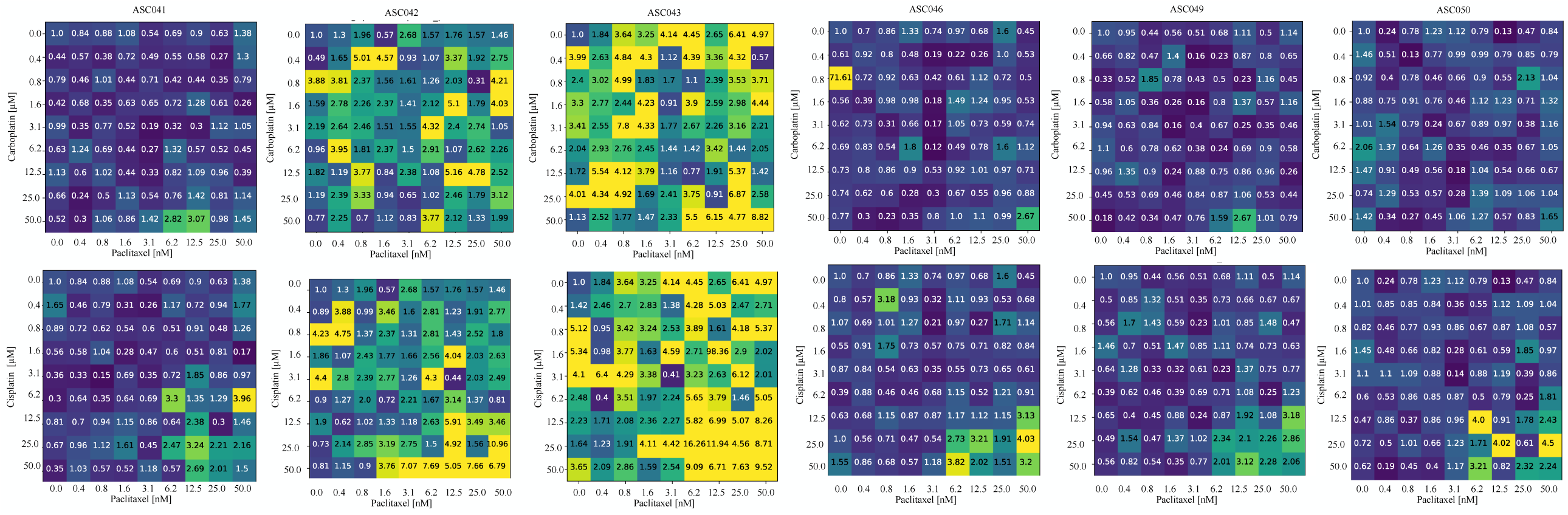
Simulated analysis of the combined treatment with cisplatin/carboplatin and paclitaxel on the digital twins. The numbers and colours represent the coefficient of variation squared of the variation in cell viability with respect to the untreated control. A total of 10 simulations for each condition were considered.

Patients ASC042 and ASC043, on the other hand, are characterised by generalised high *CV* ^2^ (3-4 fold higher on average) and no clear relationship with the drug concentration. This is also confirmed by the overall *CV* ^2^ distribution for each patient (Figure A).

Population and cell-cell variability have been extensively linked to cancer progression, evolution and the insurgence of drug resistance [23–26]. Indeed, the wider variety of possible outcomes increases the resilience of the population and its ability to withstand challenges (e.g. pharmacological treatment). The contrast with the behaviour observed for the other patients, which represent both earlier and later disease stages, suggests that this high variability status might be a temporary transition state between drug-sensitive and drug-resistant phenotypes.

While further testing on a larger cohort would be required to confirm this hypothesis experimentally, the magnitude of the difference between the *CV* ^2^ observed for patients ASC042 and ASC043 and that of the rest of the cohort points to an intrinsic difference in their response to treatment that is not captured by average values. This result underscores the fundamental role of cell-cell and population variability in understanding HGSOC progression and behaviour. Computational models are fundamental tools in this regard as they are not affected by many of the constraints that hamper the *in vitro* study of cell-cell variability (e.g., lack of single-cell levels analysis techniques, and limited availability of samples).

New methods and techniques to overcome these limitations are however required to ensure that the computational models can be appropriately validated and thus produce reliable results.

## 3 Discussion

In this work, we have presented ALISON, an *in silico* framework for the simulation and study of HGSOC cell behaviour and treatment response. It recreates a virtual epithelial tissue where healthy and cancer cells interact with each other and the environment within a defined 3D structure. Several complex behaviours emerge from this interplay, including organotropism [29] and phenotypic transitions [30]. The *in vitro* analysis of such processes, however, poses many challenges from the identification of appropriate experimental models, to their reliable use in the lab and the availability of experimental techniques for their analysis. Computational tools such as ALISON can complement the *in vitro* analysis and help mitigate the effect of necessary approximations and simplifications. As an example, the quantification of cell adhesion in the lab is a population-level measurement that might not accurately capture the complexity of this phenomenon (Figure 2a.): ALISON keeps track of each individual cell, thus potentially enabling the study of how different parameter sets, or combinations thereof, affect the ability of cancer cells to adhere to healthy tissue.

The reliability of ALISON’s results is supported by the experimental validation that was presented in this work. A panel of 6 cell lines extensively used for the *in vitro* modelling of HGSOC (Table 1) was considered, together with the 3 drugs that constitute the first-line standard of care for this disease. A good performance was measured overall, with a high correlation between simulated and experimental *IC*_50_ values (Figures 4 and 6).

A unique feature of ALISON is the possibility of using the population of models approach to simulate cell heterogeneity. Tumour phenotype variability is emerging as a defining hallmark of HGSOC, largely responsible for poor patient outcomes and low treatment effectiveness [1]. The possibility of simulating its role in metastasis and treatment response holds great potential and yields useful insights. Indeed, the simulation of heterogeneous populations led to improved accuracy in the modelling of the later-stage disease (OAW28, OVCAR4), while homogeneous virtual cells more accurately captured the behaviour of experimental models representative of an earlier HGSOC stage (Figure A1). This correlates well with the notion that diversification is an adaptive strategy that confers resilience and aids the survival of cancer cells.

The *in vitro* study of population heterogeneity is largely limited by the available technologies but the results presented in this work support the suitability of ALISON as a tool to bridge this gap and study the correlation between cell-to-cell variability and treatment response, and the development of potential strategies to reduce the impact of heterogeneity on patient prognosis.

The possibility of extending the analysis to primary samples is another key feature of ALISON and this work. A procedure to identify the model’s parameters from standard clinical information was developed. This method is flexible, allowing for missing information, and yet widely applicable as it relies on clinical data commonly available for HGSOC patients. The comparison between the simulated and experimental data provides further confirmation of ALISON’s performance and extends its scope from a preclinical research tool to a framework that could potentially be applicable in the clinic. Indeed, while the accuracy of the simulation is less consistent when patient samples are considered, our results maintain their validity in this setting, especially when clinically relevant drug doses are considered (Figure 8). These results support another potential application of ALISON, namely the identification of drugs’ efficacious ranges for specific cell lines or primary-derived cultures (i.e. likely *IC*_50_ range). This information would be useful during the optimisation of experimental conditions for *in vitro* experiments, but could also support treatment selection in the clinical setting, by determining which drug would be effective at the lowest dose.

It is also worth mentioning that one of the experimental models used for this analysis (2D monolayer) is very simple and potentially not representative of patients’ responses. Indeed, the *in vitro* analysis of sample ASC043 shows a response to platinum, especially cisplatin, with a reduction in cell viability of about 60% upon administration of even small doses of this agent. The simulation of this patient’s digital twin, however, doesn’t follow this pattern and highlights a potential treatment resistance, similarly to what was observed for sample ASC046 (Figure 8 and Table 4).

The outcome for both individuals is more in line with the *in silico* analysis as both patients passed away within one year of sample collection. While this comparison is limited, and none of the patients in the cohort were administered the simulated treatments, the results presented here support the feasibility of this approach and highlight the need for a more structured comparison in a larger cohort.

The relevance and usefulness of ALISON are further supported by the simulation of the combined administration of platinum and taxane agents, generally used as a first-line treatment for HGSOC (Figures 8 and 10, Figure A). Indeed, while the average patient response was fairly uniform across the cohort, the inter-simulation variability highlighted 2 distinct patterns of response. Earlier-stage diseases and late-stage potentially drug-resistant tumours showed a direct correlation between variability and drug concentration while intermediate samples (ASC042, ASC043) were characterised by a generalised high *CV* ^2^ irrespective of the treatment condition. This high variability might result from an unstable transition state between two stable configurations (drug-sensitive and drug-resistant phenotype) and its spontaneous emergence in our simulations is of great interest. Indeed, it suggests that the framework that we have developed is sufficiently accurate to capture this phenomenon and could thus be used to further its study *in silico* and potentially develop strategies to prevent it or lessen its effects. The presence of a similar triphasic pattern in the invasion measurements (compare Figures 2b and c with Figure A) further strengthens our observations.

While further work is needed to translate ALISON in the clinic, its features ensure that the integration of this tool within the clinical workflow would cause minimal disruption. Indeed the use of standard clinical information for the calibration of the digital twins, while potentially suboptimal, minimally impacts current procedures. The choice of focusing on metastatic disease and the colonisation of the omentum is another important feasibility feature. Indeed, for early-stage HGSOC, surgery is generally curative and most of the challenges with treatment selection only apply to metastatic and recurrent disease. The programmable, modular nature of ALISON is also highly relevant to its potential clinical translation as it facilitates its further development and the inclusion of new information. This could include new markers of treatment response or the expression of specific transport proteins that can pump drugs out of the cells thus lowering their effective concentration. New treatments could also be included as they become available. Particularly valuable would be the inclusion of PARP inhibitors, which have become a cornerstone of HGSOC treatment through the induction of synthetic lethality in HRD-positive tumours. This could be modelled in ALISON by preventing cancer cells from proliferating and promoting their death with a probability proportional to the local drug concentration.

Finally, comparing the simulations to cancer cells extracted from the ascites is key as this sample is emerging as a liquid biopsy for ovarian cancer, being more accessible than solid tissue and often available in large volumes [31]. Ascites has also been shown to provide an accurate and comprehensive picture of each patient’s disease [32] thus potentially supporting the monitoring of HGSOC and an evaluation of its evolution over time. Coupling this analysis with *in silico* simulations using a digital twin could provide valuable information and support clinicians and patients throughout the treatment. The digital twin calibration procedure could be modified for this use case to include metrics more closely capturing the dynamic evolution of HGSOC (e.g. KELIM score) and become less dependent on static clinical data (e.g., stage, surgery outcome) that might lose relevance over time.

A more extensive validation of ALISON is however required. A larger more diversified cohort is needed to test the model’s accuracy on a wider range of conditions and behaviours. Experimental data acquired in more *in vivo*-relevant conditions (e.g., organoids) would also be required to further strengthen the validity of the simulated results. A comprehensive drug panel, including the treatment that each patient is undergoing, would also be necessary, as it would enable the comparison with the actual patient outcome and would provide more useful insights beyond the standard of care.

## 4 Conclusion

Precision medicine in the oncological field has mainly relied on the identification of specific genetic variants (druggable targets). This approach is largely unsuitable for diseases with few recognised markers of outcome or when the combined effect of multiple mutations needs to be assessed. Tools like ALISON could prove valuable in this regard, enabling the integration of diverse clinical information in a unified framework. While further work is required to increase the model’s accuracy on patient samples and verify its robustness on a larger cohort, the methodological framework maintains its validity and generality. Indeed, changing the structure of the virtual tissue and the characteristics of the AB model (cell types, states, behavioural rules) would be sufficient to adapt this framework to simulate different tissues either in physiological or pathological conditions.

As such, ALISON’s scope of application is wide, ranging from a preclinical research tool, capable of complementing the experimental analysis with information difficult to measure experimentally, to a clinical instrument, providing useful insights on disease stage and treatment options.

Overall, our methods and results demonstrate how computational models and advanced data analysis methods can uncover relevant information on cancer cell behaviour and simulate cell-cell and population heterogeneity, a feature difficult to capture with currently available methods, thus supporting treatment personalisation in HGSOC.

## 5 Methods

### 5.1 Cell culture

A panel of 6 HGSOC cell lines (CaOV3, OAW28, OVCAR4, OVSAHO, PEO1, PEO4) were used for ALISON’s development and validation. They were chosen as their features (Table 1) are consistent with the characteristics of a typical HGSOC patient cohort, mostly late-stage disease, with a notable proportion of drug-resistant cells. Genetic features with an established link to treatment response in HGSOC are also included. BRCA1/2 genes are part of the homologous recombination pathway and the main indicators of HRD status [33]. NF1, on the other hand, is a tumour-suppressor gene whose mutation has been associated with treatment resistance [34].

CaOV3, PEO1 and PEO4 cells were kindly donated by Prof. Deborah Marsh, University Technology Sydney, while Prof. Nikola Bowden, University of Newcastle generously provided OAW28 and OVSAHO cells. OVCAR4 cells were purchased from the American Type Culture Collection (USA). OAW28, OVSAHO, PEO1 and PEO4 cells were maintained in RPMI medium (Thermo Fisher, USA), while CaOV3 and OVCAR4 cells were kept in DMEM (Sigma-Aldrich, USA). In all cases, media were supplemented with 10% FBS (Scientifix, Australia), 1% Pen-strep (Sigma-Aldrich, USA) and 1% GlutaMAX (Thermo Fisher, USA). All cell lines were confirmed free of mycoplasma infection and validated via STR profiling.

Primary cells were obtained from ascites, a liquid that accumulates in the abdominal cavity of HGSOC patients and has been shown to contribute to cancer dissemination by facilitating cell movement within the abdomen [13, 14]. Samples were collected from patients undergoing paracentesis at the Royal Hospital for Women following a HGSOC diagnosis (ethical approval provided by the South Eastern Sydney Local Health District Human Research Ethics Committee (HREC), reference ID Royal Hospital for Women HREC 19/001). Cancer cell cultures were obtained by adding 25 ml of ascites fluid to a T175 cell culture flask and progressively substituting it with cell culture media over a few days. Adherent cells were then maintained in RPMI medium (Thermo Fisher, USA), supplemented with 10% FBS (Sigma-Aldrich, USA), 1% Pen-strep (Sigma-Aldrich, USA) and 1% GlutaMAX (Thermo Fisher, Waltham, MA, USA). The study limited cells to initial passages (up to passage 3) to avoid alterations in primary cells’ behaviour.

A portion of the experimental analysis relied on a 3D organotypic model of the peritoneal lining (i.e., omentum) to increase the *in-vivo* relevance of the data. This model, initially presented in [15], involves co-culturing cancer cells, fibroblasts and mesothelial cells in a 3D structure further described in the next section. Fibroblasts and mesothelial cells were extracted from omentum samples collected from patients undergoing surgery for benign or non-metastatic conditions at the Royal Hospital for Women and Prince of Wales Private Hospital (site-specific approval ethics LNR/16/POWH/236). The tissue was processed as previously described [15] and the isolated cell populations were maintained in DMEM media supplemented with 10% FBS (Sigma-Aldrich, USA), 1% non-essential amino acids (Sigma-Aldrich, USA) and 1% GlutaMAX (Thermo Fisher, USA) and 1% Antibiotic-Antimycotic (Gibco, USA). Informed consent was obtained from all the patients involved in the study.

### 5.2 Quantification of adhesion

Adhesion was quantified in a 3D organotypic model of the omentum as previously described [15, 35]. Briefly, 100 *µ*l of a solution of media, fibroblast cells (4 10^4^ cells/ml) and collagen I (5 ng/*µ*l, Sigma-Aldrich, USA) was added to the wells of a 96-well plate. About 4 hours later 20,000 mesothelial cells were seeded on top in 50 *µ*l of media, to mimic the structure of the lining of the peritoneal cavity [15]. After 24 h the media was changed to 100 *µ*l containing 1,000 (PEO4, OVCAR4, CaOV3, OAW28) or 2,000 (PEO1, OVSAHO) cancer cells.

Adhesion was measured at 2,3 and 4 h after cancer cell seeding, by fixing the culture with 96% ethanol for 10 minutes and staining with a solution of 1% crystal violet in ethanol. Each well was then washed extensively with running d*H*_2_O to remove excess stain. Cells were then lysed with 50% acetic acid and their density was quantified with an absorbance measurement (at 595 nm) [35]. For each cell line, 3 biological replicates, each comprising at least 2 technical replicates, were conducted. Each biological replicate relied on a different donor for the healthy cells.

The analysis was conducted using custom-made software written in Python (v3.9). The technical and biological replicates were averaged and the average absorbance at 4h was used as normaliser.

### 5.3 Quantification of Invasion

Invasion was evaluated in the same 3D organotypic model of the omentum initially described in [15] and further detailed in the previous section. The only difference with respect to the quantification of adhesion was that the model was seeded in transwell insert (24-well plate version Corning Life Sciences, USA) rather than in a regular 96-well plate. Additionally, upon cancer cells seeding (2 10^6^ cells/ml for PEO1 and OVSAHO and 1 10^6^ cells/ml for PEO4, OVCAR4, CaOV3 and OAW28) a nutrient gradient was established (20% FBS in the bottom chamber and 1% FBS media in the transwell insert) to encourage cell invasion. The cultures were incubated for 24 (PEO1) or 48 (PEO4, OAW28, OVSAHO, CaOV3, OVCAR4) h before gently washing transwell inserts with PBS and fixing cells in 4% paraformaldehyde for 10 minutes. Wells were washed again in PBS and mounted on a microscope slide using DAPI mounting medium (Fluoroshield, Sigma-Aldrich, USA) [35]. Slides were allowed to dry for at least 1 h prior to imaging at the microscope (Leica DM 2000 LED fitted with a Leica DFC450c camera). Ten images from different regions of the slide were acquired [35]. Cell counting was automatically conducted using ORACLE [18] a deep neural network that we developed to enable the analysis of images from transwell invasion assays and support the classification of healthy and cancer cells without the need for cell-type specific staining or transfection with fluorescent markers. During the analysis, images were divided into 16 non-overlapping sub-images due to computational constraints. The number of cancer cells/image was then transformed in the fraction of the total image area occupied by cancer cells (number of pixels classified as part of cancer cells divided by the total number of pixels), to eliminate the dependence on the size of the image and increase the similarity between this metric and the one used in the simulations (fraction of the healthy tissue occupied by cancer cells).

### 5.4 Drug response

Response to cisplatin, carboplatin and paclitaxel was measured on both cell lines and primary cells in 2D monocultures. The change in the experimental model was due to the results of a preliminary analysis [35] aimed at determining which combination of experimental data would yield the most accurate computational model.

An initial population of 50 10^3^ cells/ml (PEO1, OAW28, CaOV3, patient-derived samples) or 100 10^3^ cells/ml (PEO4, OVCAR4, OVSAHO) cells was seeded in a 96-well plate using a volume of 100 µl/well. After 24 h the media was substituted with DMEM or RPMI containing different concentrations of cisplatin (50, 25, 12.5, 6.25, 3.12, 1.6, 0.8, 0.4 µM), carboplatin (50, 25, 12.5, 6.25, 3.12, 1.6, 0.8, 0.4 µM) and paclitaxel (50, 25, 12.5, 6.25, 3.12, 1.6, 0.8, 0.4 nM). In all cases, untreated cells and media supplemented with 5% DMSO were used as controls.

Cell viability was quantified after 72 h with an MTT assay (Thermo Fisher, USA). Three biological replicates for each cell line, each comprising three technical replicates were acquired. The absorbance measurements at 570 nm were averaged and normalised with respect to the negative controls.

### 5.5 ALISON development and simulation

ALISON combines a FEM, realised using the Python library fenics (2019.1.0) and an AB framework realised *ad-hoc* in Python (v. 3.9) (Figure 1). Both models have a temporal resolution of 1 h and rely on the same mesh, a cylinder of height 11 mm and radius 4.25 mm and 42407 tetrahedral elements (Figure 1b). These measures match the height and diameter of the wells in a 96-well plate and the organotypic model of the omentum used for part of the *in vitro* analysis.

The FEM is used to dynamically solve the heat equation (Equation 3) and thus model the diffusion of oxygen, glucose, lactate and each of the three considered drugs.

A(**x**,t) is the distribution of the simulated variables over space (**x**) and time (t). *C_v_* (**x**) and *ρ*(**x**) are the heat capacity and mass density of the organotypic model. They were both maintained uniform throughout the structure. In this context, the heat capacity measures the ability of the virtual tissue to absorb the mass flux (oxygen, glucose, lactate, drugs). The system has no leaks and we expect the simulated concentrations to be below saturation. Hence the heat capacity was set to 1 *mm*^3^/*h*^2^. Most human tissues have a mass density of about 1 mg/*mm*^3^ (ITIS foundation database). As such, *ρ*(**x**) was set to 1 mg/*mm*^3^.

k(**x**) is the thermal conductivity and is connected with the molecules’ fluxes that arise due to differences in concentration. It was set to 0.594 *mm*^2^/h, in accordance with experimental measurements conducted in collagen gels [36]. Finally, S(**x**,t), is the source distribution, that is the current position of the cells. This matrix is updated at each iteration to reflect the status of the population.

A scale factor of 1:10 was applied due to computational constraints. This consists of reducing by a 10-fold factor the number of cells, the volume of the virtual organotypic model and the concentrations/rates of production of the simulated molecules. This factor can be adjusted in the configuration files.

The initial oxygen level at each node of the mesh was set to 5.85 10*^−^*^8^, as the concentration of oxygen within the incubator in standard culturing conditions is 1241 mg/l. Similarly, the glucose level was set to 9.43 10*^−^*^5^ in accordance with the concentration of this molecule in RPMI media. The starting lactate level was set to 0, and the drug level, when relevant, was set to be equal to the experimental value.

Mixed boundary conditions were employed, Dirichlet at the top of the mesh and Neumann at the bottom and side of the mesh. This is coherent with the experimental model, where the conditions at the top of the organotypic model/peritoneal cavity are largely constant and not affected by the activity of the cells. Similarly, the walls of the well prevent any mass flux at the bottom and sides of the organotypic model.

The AB model, on the other hand, describes the behaviour of individual cells, through a set of specific states that each cell type can assume, and programmable user-defined rules that formalise allowed behaviours and corresponding likelihood of occurrence. As an example, in Equation 4 a cell in status s (*C_s_*) can transition to status e (*C_e_*) with probability *P_s−>e_*. The latter is a user-defined function that can combine constant parameters, environmental conditions, cell-specific information and signals from other cells. Figure 1c shows the structure of the AB model used in this work and the probability functions driving its dynamic behaviour.

All simulations presented in this work were conducted using the computational cluster Katana supported by Research Technology Services at UNSW Sydney [37].

### 5.6 Parameters identification

The identification of the parameters describing the behaviour of healthy cells relied on virtual organotypic models containing only fibroblasts or mesothelial cells. A number of configurations equal to 100x the number of parameters to be estimated (600 for fibroblasts and 500 for mesothelial cells) were chosen using the Latin hypercube sampling method (Python library scipy.stats.qmc). Each configuration was simulated 3 times. The number of living cells at the end of each simulation was computed and compared with the starting condition. As limited information on the behaviour of healthy cells in the organotypic model is available, we have assumed their density to be constant within the experiments. As such, the parameter sets associated with the smallest difference between initial and final cell density were selected (Table A1). When more than one configuration yielded the same optimal result, the first one was chosen.

The parameters for the cancer cell lines were determined through a similar approach. Initially, only the untreated condition was considered and an organotypic model including all 3 types of cells (fibroblasts, mesothelial and cancer cells) was used. 700 parameter configurations were determined with the Latin lattice hypercube sampling, and each was simulated 3 times.

For each configuration, the average doubling, adhesion rate and invaded area were obtained and compared to the experimental values at the relevant time points. The expected population size at the end of the experiment was obtained by dividing the total number of living cancer cells at the end of the simulation by the size of the initial cancer population. It was then compared with the theoretical value computed from the doubling rate. Adhesion was defined as the number of cancer cells in the mesothelial layer at the relevant time points (1 - 4 h post seeding), while invasion was obtained as the percentage prevalence of cancer cells in the fibroblast layers at either 24 or 48 h. Experimentally measured adhesion and invasion data were used in the comparison (further details in the previous sections), while the doubling time for each cell line was determined from the literature (Table 1).

A cost function comparing the experimental and simulated data was then computed for each configuration (Equation 5). It quantifies the difference between the *in-silico* and *in-vitro* doubling (first term), adhesion (second term) and invasion (third term). The last term was introduced to favour configurations associated with a higher number of transitions between different cell statuses. The rank function was used, instead of the actual value of the scores, to offset numerical differences between the scores and thus equally weight each component. The bias term was further used to penalise configurations accurately capturing only a subset of the metrics in Equation 5. Indeed, if the coefficient of variation of the individual score components was above 0.2, the bias term was set to 2. In all other cases, it was assigned value 1.

The values of the cost function provided a metric of the accuracy of each parameter configuration in recapitulating the behaviour of the corresponding cell lines.

The best configuration for each cell line (Table A3) was then used to calibrate the drug response parameters (the modified ALISON model and corresponding probability functions are available in Figure A3). For each drug, 200 parameter configurations, estimated with the Latin hypercube sampling method, were considered. Treatment with an amount of drug corresponding to the *IC*_5_0 dose (Table 6) was simulated for each cell line and drug and compared to the expected population in untreated conditions. As per the definition of *IC*_50_, the configuration closest to a ratio of 0.5 was determined to be optimal. Again the resulting score function was saved to serve as a reference distribution for the simulation of cell-cell variability.

**Table 6.** *IC*_50_ values used during the calibration of ALISON.

| Cell line | Carboplatin [M] | Cisplatin[M] | Paclitaxel [M] |
| --- | --- | --- | --- |
| CaOV3 | $24.1 \cdot 10^{-6}$ [44] | $1.3 \cdot 10^{-6}$ [45] | $1.0 \cdot 10^{-9}$ [46] |
| OAW28 | $50.0 \cdot 10^{-6}$ [47] | $10 \cdot 10^{-6}$ [48] | $3.0 \cdot 10^{-9}$ [48] |
| OVCAR4 | $60.0 \cdot 10^{-6}$ [47] | $8.0 \cdot 10^{-6}$ [49] | $5.7 \cdot 10^{-9}$ [50] |
| OVSAHO | $6.0 \cdot 10^{-6}$ [51] | $3.7 \cdot 10^{-6}$ [52] | $3.4 \cdot 10^{-9}$ [53] |
| PEO1 | $20.0 \cdot 10^{-6}$ [54] | $0.5 \cdot 10^{-6}$ [45] | $3.6 \cdot 10^{-9}$ [45] |
| PEO4 | $45.0 \cdot 10^{-6}$ [55] | $10.0 \cdot 10^{-6}$ [56] | $3.0 \cdot 10^{-9}$ [45] |

### 5.7 Cell-cell variability simulation

Cell-cell variability was implemented in ALISON by assigning each virtual cell a different parameter set. The likelihood of choosing each specific configuration was inversely proportional to their score, that is parameter sets yielding a behaviour closer to the experimentally measured one were more likely to be chosen. A random selection with replacement (function random.choices from the numpy package) was used, and the weight sequence was obtained by transforming each configuration’s score (score(c)) using Equation 6. For cell lines simulations the scores were the ones computed as in Equation 5 while for patient-derived samples they were obtained during the digital twin calibration described in the next section.

At cell division, the mother cell would maintain the same parameter set while the daughter cell was assigned a new parameter set following the same procedure. The parameters associated with response to treatment were chosen separately from the ones also present in the untreated condition, similarly to the procedure used during parameters calibration.

In this work, cell-cell variability was simulated only for cancer cells, but the same approach can be easily extended to the simulation of fibroblasts and mesothelial cells’ behaviour.

### 5.8 Digital twin calibration

The digital twin calibration procedure was developed as a 2 step process (Figure 8). Initially, a subset of patient features (recurrence status, known drug resistance, genetic profile) were matched to the cell lines to determine which combination of cell lines best represented each individual (percentage similarity index and unbiased fractions).

The additional clinical information (age, stage, debulking surgery and the CA125 level closest to sample collection) was then used to calculate a prognosis bias. In particular, an age above the average diagnosis age for HGSOC (63 years [38]), a higher stage, an elevated CA125 and residual disease following surgery would generate a bias toward a worse prognosis, with a magnitude proportional to the feature’s relevance (e.g., a highly elevated CA125 would yield a stronger bias than a moderately altered one). Opposite features would result in a bias toward a better prognosis. The bias for each factor was calculated independently and summed to determine each patient’s overall bias.

A bias toward a worse prognosis would be added to the percentage similarity of the most aggressive cell lines in our panel (OAW28, OVCAR4, PEO4) and subtracted from the others. Conversely, a bias toward a good prognosis would increase the tumour’s similarity to CaOV3, OVSAHO and PEO1 cells. The biased cell line proportions were then re-normalised so that their sum would be 1. The score distributions for each cell line were then combined, by calculating the weighted average using the biased fractions as weights, to create the reference parameter distribution for each patient.

To improve the flexibility and applicability of this method, no clinical information is mandatory for the calibration of the digital twin. Any missing field will be ignored in the procedure.

### 5.9 Validation and analysis

ALISON was validated on both cell lines and patient-derived samples. A total of 10 simulations for each condition were considered. The cell lines validation was conducted simulating the entire dose-response curves to cisplatin, carboplatin and paclitaxel and comparing them to experimentally measured results. The *IC*_50_ for both experimental and simulated data was obtained by fitting the available data points to a sigmoid function (Equation 7), where x is the drug concentration, y the resulting decrease in viability and L, k, x0 and b the parameters to be optimised. The drug concentration associated with a decrease in population size equal to 0.5 was then determined.

Both the optimal parameter configuration and the cell-cell variability case were simulated. This enabled the study of the effect of this new feature and confirmed the accuracy of our model for the simulation of cell-line data.

A similar analysis was conducted for the patient-derived sample, simulating the dose-response curve for each drug and comparing the results to the experiments. In this case, only the cell-cell variability configuration was considered as its features more closely resemble the behaviour of primary cells.

Finally, combined treatment with a platinum-based agent (cisplatin or carboplatin) and paclitaxel, in the same doses as before was simulated. This is a clinically relevant configuration as these therapeutic agents are largely administered together. The average decrease in cancer population size, together with the inter-simulation variability (*CV* ^2^ Equation 2) were studied.

## Declarations

### 5.12 Ethics approval and consent to participate

All patients participating in this study provided informed consent. Ascites samples were collected at the Royal Hospital for Women (ethical approval provided by the South Eastern Sydney Local Health District Human Research Ethics Committee (HREC), reference ID Royal Hospital for Women HREC 19/001).

Healthy omentum tissue was collected from patients undergoing surgery for benign or non-metastatic conditions at the Royal Hospital for Women and Prince of Wales Private Hospital (site-specific approval ethics LNR/16/POWH/236).

### 5.13 Code availability

ALISON code is available at https://github.com/MarilisaCortesi/ALISONsimulator

## 6 Equations

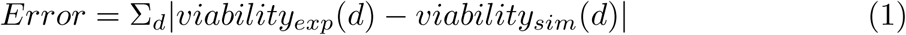

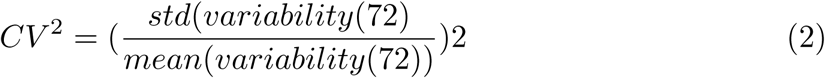

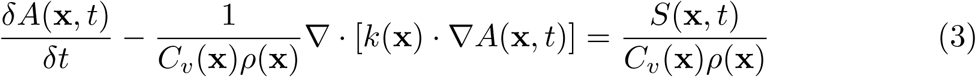

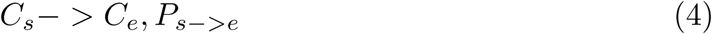

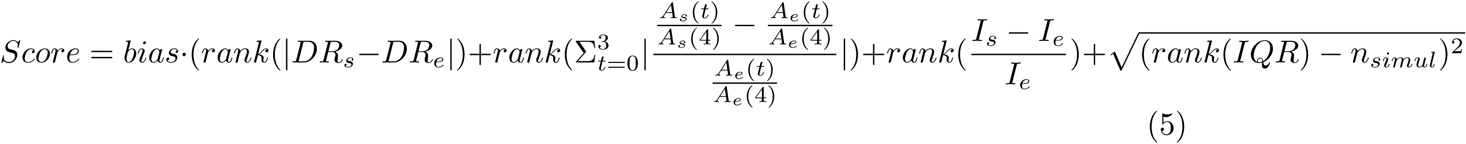

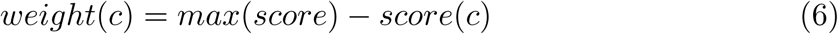

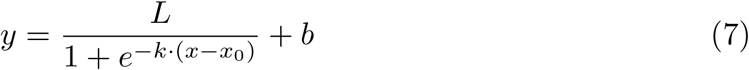

## Appendix A Extended data

**Fig. A1.**
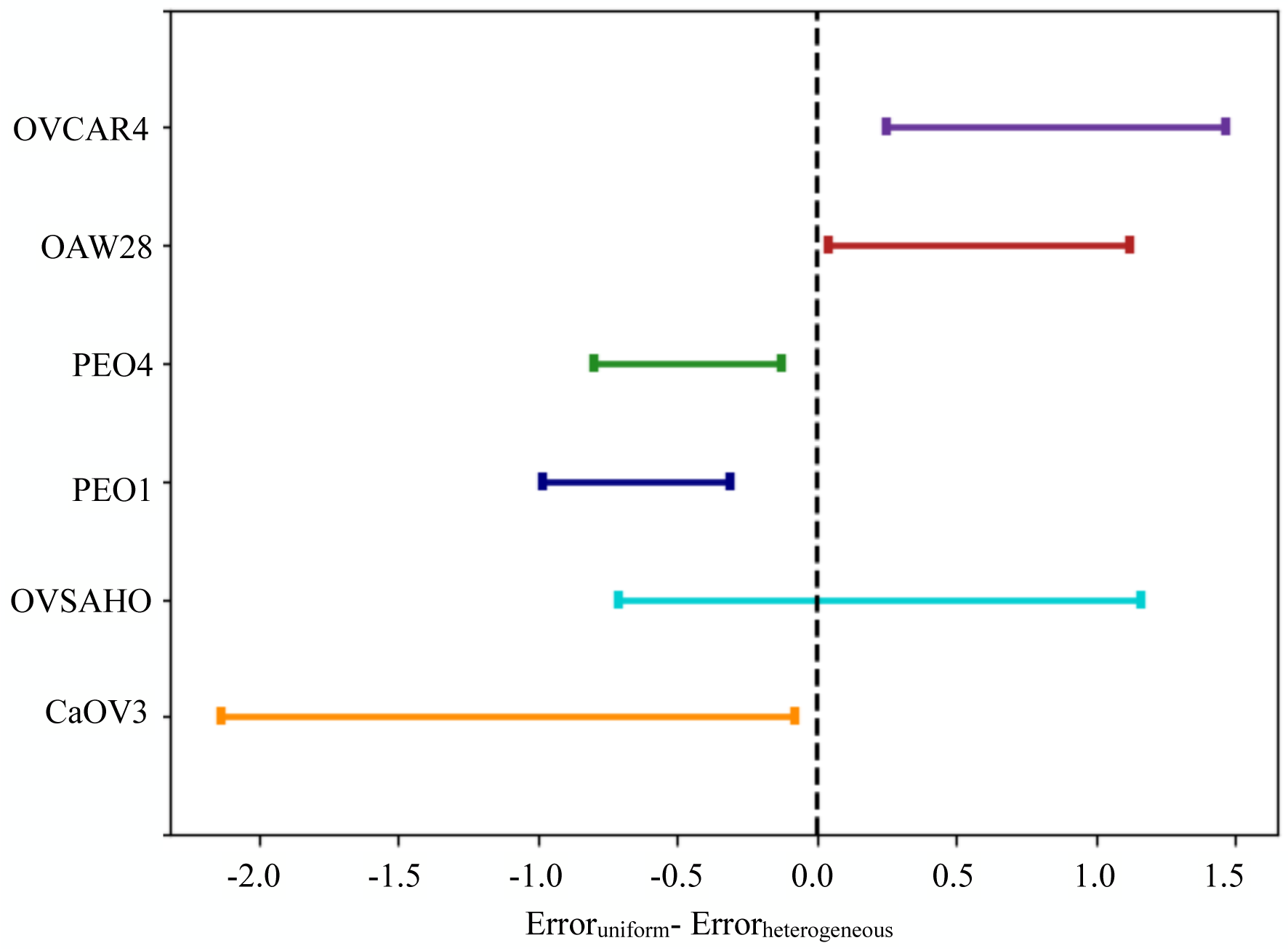
Change in error function recorded when cell-cell variability was included. Positive values on the x-axis correspond to an improvement in accuracy when variability was added.

**Fig. A2.**
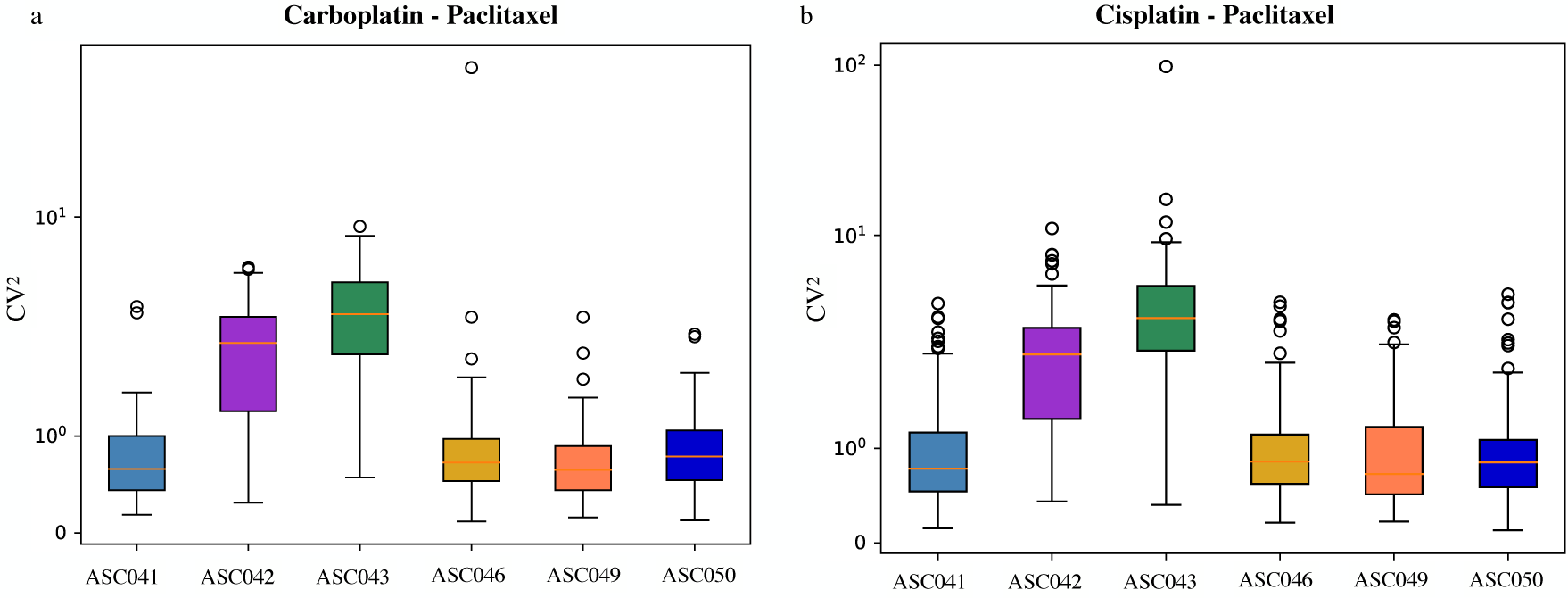
Distribution of the *CV* 2 for each patient and combined treatment. a. Refers to the carboplatin-paclitaxel treatment while in b. The data for the cisplatin-paclitaxel treatment are shown.

**Fig. A3.**
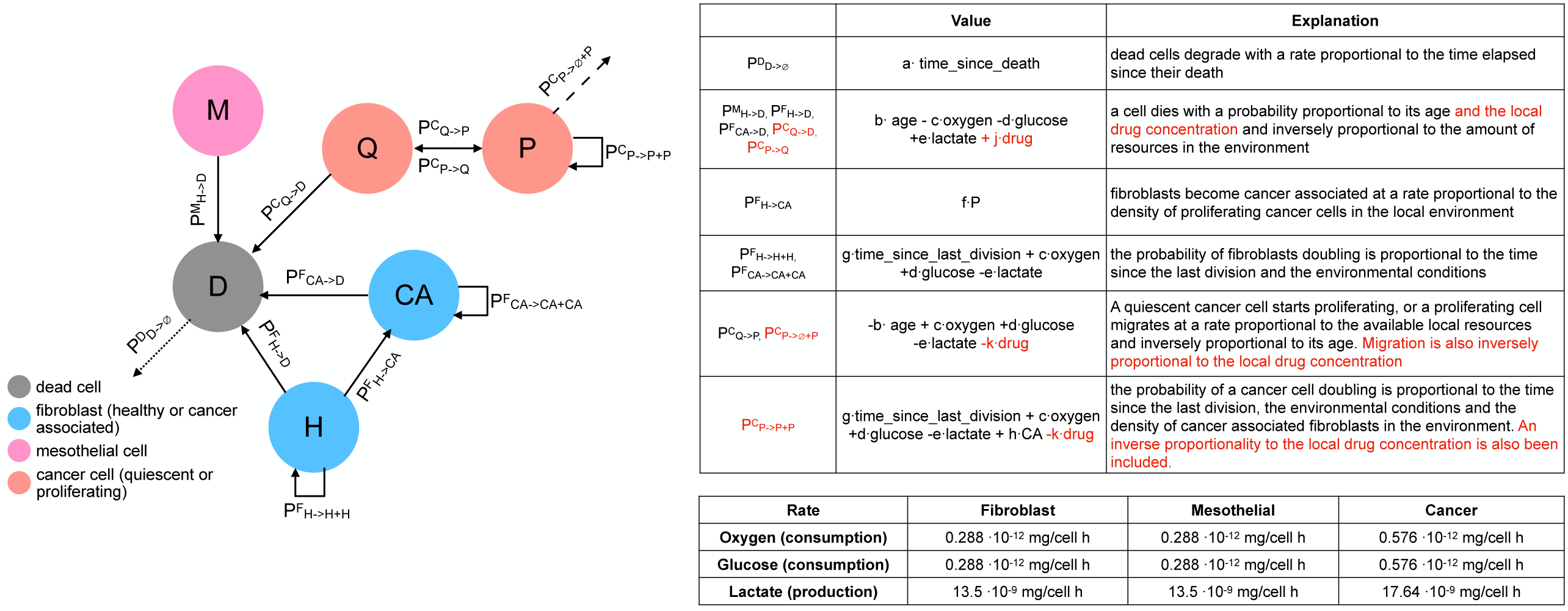
Modified ALISON model to include drug treatment. Cell statuses and transitions were not changed, the only modifications were in the probability functions and are shown in red in the table.

**Table A1.** Rates of consumption (oxygen and glucose) and production (lactate) used for the simulations.

| Rate | Fibroblast | Mesothelial | Cancer |
| --- | --- | --- | --- |
| Oxygen (consumption) [mg/cell h] | $0.288 \cdot 10^{-12}$ | $0.288 \cdot 10^{-12}$ | $0.576 \cdot 10^{-12}$ |
| Glucose (consumption) [mg/cell h] | $17.64 \cdot 10^{-6}$ | $17.64 \cdot 10^{-6}$ | $35.28 \cdot 10^{-6}$ |
| Lactate (production) [mg/cell h] | $17.64 \cdot 10^{-6}$ | $17.64 \cdot 10^{-6}$ | $35.28 \cdot 10^{-6}$ |

**Table A2.**
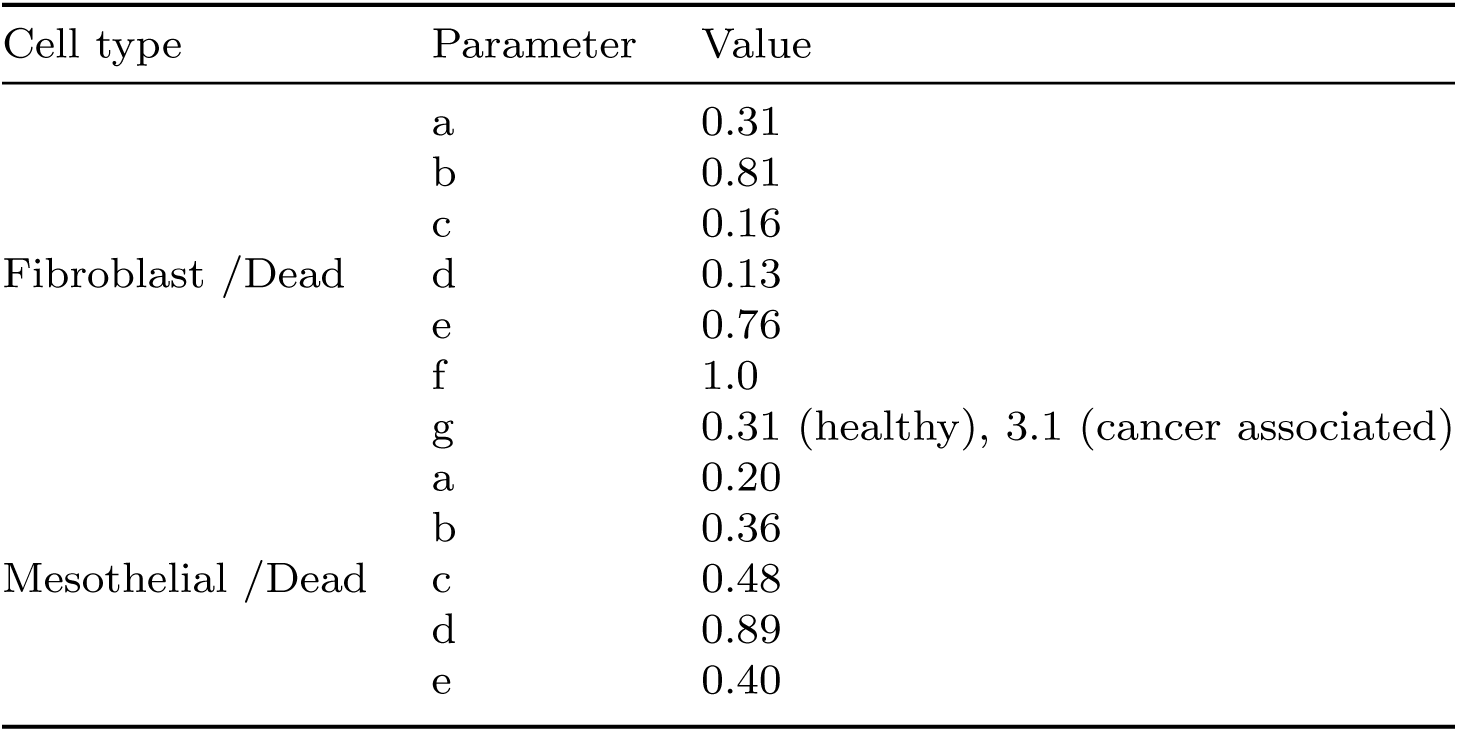
Optimal parameter configurations for fibroblasts and mesothelial cells.

**Table A3.**
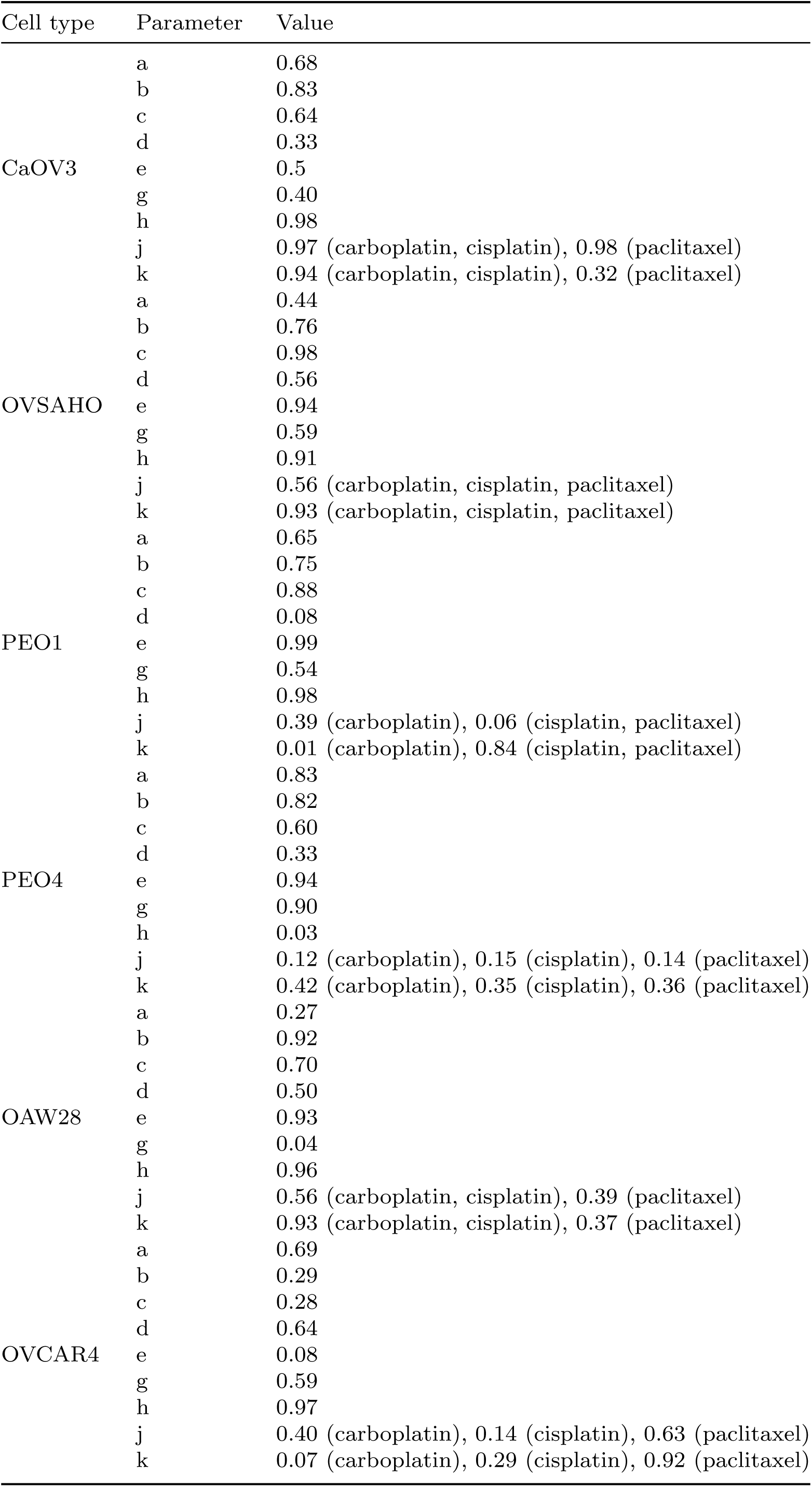
Optimal parameter configurations for each considered cancer cell line.

**Table A4.** Breakdown of the bias used in the digital twin calibration to obtain the biassed fractions.

| Bias | ASC041 | ASC042 | ASC043 | ASC046 | ASC049 | ASC050 |
| --- | --- | --- | --- | --- | --- | --- |
| Age | -0.03 | 0.13 | 0.18 | -0.15 | -0.06 | 0.0 |
| Stage | 0.88 | 0.39 | 0.21 | 0.39 | 0.88 | 0.39 |
| CA125 | 0.20 | 0.18 | 0.64 | 0.23 | 0.70 | 0.79 |
| Debulking | 0.0 | 0.0 | 1.0 | 0.0 | 1.0 | 1.0 |
| Total | 1.04 | 0.70 | 2.02 | 0.47 | 2.52 | 2.19 |

